# NLR from soybean *Rsv1* locus confers broad-spectrum resistance to soybean mosaic virus G1-G7 strains by recognizing viral P3 protein

**DOI:** 10.64898/2026.06.29.735421

**Authors:** Heyu Zhao, Bei Gou, Jiaqi Liao, Yanxiao Zhao, Tongqing Yang, Peilin Huang, Yingying Zhu, Yi Tie, Mohan Wang, Long Gao, Kai Li, Haijian Zhi, Xiaoyan Cui, Xin Chen, Yi Xu, Kaixuan Duan, Yuanchao Wang, Xiaorong Tao

**Affiliations:** State Key Laboratory of Agricultural and Forestry Biosecurity, Nanjing Agricultural University, Nanjing, 210095, China; Key Laboratory of Soybean Disease and Pest Control (Ministry of Agriculture and Rural Affairs), Nanjing Agricultural University, Nanjing, Jiangsu 210095, China; Key Laboratory of Plant Immunity, Department of Plant Pathology, Nanjing Agricultural University, Nanjing, 210095, China; Key Laboratory of Integrated Management of Crop Disease and Pests, Ministry of Education, Nanjing Agricultural University, Nanjing, 210095, China; Key Laboratory of Soybean Biology and Genetic Breeding, Ministry of Agriculture and Rural Affairs, Nanjing 210095, China; College of Agriculture, Nanjing Agricultural University, Nanjing 210095, China; National Laboratory of Crop Genetics and Germplasm Innovation and Utilization, Nanjing 210095, China; Institute of Industrial Crops, Jiangsu Academy of Agricultural Sciences/Jiangsu Key Laboratory for Horticultural Crop Genetic Improvement, Nanjing, Jiangsu, 210014, China

**Author notes:** Correspondence author. Xiaorong Tao (X.T.), Yuanchao Wang (Y.W.). These authors contributed equally (Heyu Zhao, Bei Gou).

## Abstract

Nucleotide-binding leucine-rich repeat (NLR) immune receptor genes are of significant value in disease resistance breeding and the control of viral diseases. *Soybean mosaic virus* (SMV) poses a serious threat to soybean production and the *Rsv1* locus in soybean cultivar Suweon 97 confers broad-spectrum resistance against SMV strains G1 to G7; however, this locus harbors no fewer than 18 NLR genes, and thus the broad-spectrum antiviral mechanisms underlying the *Rsv1* locus remain poorly understood to date. Here, we established a rapid and highly efficient screening system for cloning NLR genes from soybean *Rsv1* locus and identified a broad-spectrum antiviral NLR gene *13g184900* from this highly complicated locus. The NLR encoded by *13g184900* can recognize viral P3 protein from all SMV strains (G1-G7) and another potyvirus *Bean common mosaic virus* (BCMV). The coiled-coil (CC) domain of this NLR directly interacts with viral P3 protein. Additionally, we showed that this NLR originated from wild soybean accession in East China and has been introduced into several soybean cultivars during domestication. Collectively, we developed a high-throughput screening system for identifying NLR genes in soybean and our study provides new mechanistic perspective on how the *Rsv1* locus mediates the broad-spectrum resistance to all SMV G1-G7 strains.

## Introduction

Soybean (*Glycine max*) is one of the important economic crops worldwide, with oil-and protein-rich seeds ranking as the world’s top vegetable oil source^1,2^. With global climate change and the intensification of worldwide trade, plant diseases have emerged as the principal constraint on global production^3,4^. In the United States, climate change is projected to reduce average soybean yields by 86 – 92% by 2050^5^. Leveraging resistance genes to breed disease resistance crop represents one of the most efficient and sustainable approaches to managing diseases^6,7^. Among these resistance genes, the nucleotide-binding leucine-rich repeat (NLR) genes are the most valuable and most widely applied category^8^. NLRs represent the largest family of resistance genes in plant genomes, accounting for more 60% of total disease resistance genes^9^. NLR proteins recognize pathogen effectors (also known as avirulence/Avr proteins) and activate effector-triggered immunity (ETI), typically accompanied by hypersensitive response (HR) to confine pathogens at initial infection sites^10,11^. Since the first NLR gene was cloned in 1994, only five soybean NLR genes have been cloned. In contrast, far greater numbers have been identified in other model and crop species, including 95 in *Arabidopsis thaliana*, 55 in *Oryza sativa*, 44 in *Triticum aestivum*, and 20 in *Solanum lycopersicum*, respectively^9^. Therefore, the resistant genes in soybean urgently require further exploration.

Among pathogens affecting soybean, SMV is a particular concern due to its widespread occurrence across major soybean-producing regions, including China, the United States, Brazil, India, and Argentina. It is also the predominant viral pathogen of soybean in China and the United States^12,13^. SMV’s infection induces a suite of symptoms including diminished seedling viability, seed coat mottling, flower abortion, and suppressed pod development, which collectively lead to considerable yield reduction and reduced seed quality^14–16^. In severe disease epidemics of China, yield losses can reach up to 50%, depending on soybean genotypes and SMV strains^17^. As early as 1948, there are works first documented that multiple SMV strains cause soybean mosaic disease. Multiple independent classification systems for SMV isolates have been developed worldwide. The widely accepted system for SMV strain classification is principally based on variations in viral virulence and phenotypic reactions on a panel of differential host lines. In the SMV classification system of the United States, SMV isolates were divided into seven strains (G1 to G7)^12^. Among these, strain G1 displays the lowest virulence: it infects only susceptible cultivars and fails to infect all tested resistant varieties. By contrast, strain G7 is the most virulent, capable of overcoming cultivar resistance, and induces necrotic symptoms and typical mosaic phenotypes on different resistant cultivars, respectively^18^.

Besides SMV, BCMV is also a critical viral pathogen threatening soybean production^19^. In addition to soybean, it causes yield losses on multiple leguminous crops including common bean, cowpea and peanut, making it one of the most economically damaging viruses affecting global legume production^20,21^. BCMV is a close relative of SMV, with highly similar genome architectures and pathogenesis mechanisms^22^. Both of SMV and BCMV are members of the genus *Potyvirus* in the *Potyviridae* family^23^. This *Potyviridae* family accounts for approximately 30% of all known plant viruses and harbours numerous economically important species that infect major staple crops worldwide^24^. Its genome contains a ∼10 kb open reading frame encoding a polyprotein, which is processed by viral proteases into 11 functional proteins: P1, HC-Pro, P3, 6K1, CI, 6K2, VPg, NIa-Pro, NIb, CP and P3N-PIPO^25,26^. In these functional proteins, the biochemical activity and pathogenic roles of the P3 protein remain unknown, making P3 one of the least understood proteins among potyvirus members^27^.

To date, four independent loci for SMV resistance, *Rsv1*, *Rsv3*, *Rsv4*, and *Rsv5*, have been identified^28–31^. *Rsv1* was the first SMV resistance locus identified and mapped to chromosome 13, and it is the most common SMV resistance locus in soybean germplasm^28^. Meanwhile, *Rsv1* locus is also the most structurally complex of all reported SMV resistance loci. It contains at least 10 alleles across different soybean cultivars: *Rsv1*, *Rsv1-t*, *Rsv1-y*, *Rsv1-m*, *Rsv1-k*, *Rsv1-r*, *Rsv1-s*, *Rsv1-n*, *Rsv1-h*, and *Rsv1-c*^28,32–36^. The *Rsv1* allele was first identified in the soybean cultivar PI 96983^32^. Researchers also mapped two SMV resistance loci in the same region on chromosome 13 of PI96983, called *Rsc-ps* and *Rsc-pm*^37^. Among all reported *Rsv1* alleles, *Rsv1-h* from the soybean cultivar Suweon 97 has the greatest significant breeding application value and potential, as it has been verified to confer resistance to all SMV G1-G7 strains in the United States^35^. However, due to the extreme complexity and high sequence divergence of the *Rsv1* locus, as well as the lack of an effective screening system, none of the resistance genes at this locus have been identified in any soybean cultivar, including the broad-spectrum resistant gene in Suweon 97.

In this study, we establish a rapid and highly efficient screening system for characterization of NLR genes in soybean *Rsv1* locus. Using this system, we identified a NLR gene *13g184900* from this highly complicated *Rsv1* locus which confers the resistance to both SMV and BCMV. The NLR encoded by *13g184900* characterized in this study can recognize P3 proteins from all SMV G1-G7 strains and also BCMV, and subsequently trigger HR. We also found that the CC domain of the NLR encoded by *13g184900* directly interacts with SMV/BCMV P3 protein. Furthermore, evolutionary analysis revealed that this resistance gene originated from wild soybean in China, has retained its functional integrity throughout soybean domestication.

## Results

### Establishing a rapid and highly efficient system for screening 18 NLR genes from the soybean *Rsv1* locus

Validating NLR functions via transforming each gene individually into soybean using conventional methods is labor-intensive and time-consuming. To circumvent this technical challenge, we adopted an *Agrobacterium*-mediated transient expression system in *Vigna unguiculata* (cowpea) to screen NLR genes from *Rsv1* locus. Cowpea and soybean both belong to the *Fabaceae* family and are expected to share conserved immune signaling pathways. Our previous study demonstrated that many cowpea varieties confers the resistance against *Cucumber mosaic virus* (CMV); using this cowpea system, expression of the CMV Avr protein 2a in cotyledons can induce a strong HR^38^ (Fig. 1a). This indicates that the *Agrobacterium*-mediated delivering of heterologous gene can be stably expressed in cowpea leaves. To identify which cowpea genotype has the highest heterologous expression efficiency, we selected six commercial cowpea varieties. CMV 2a induced HR in all six tested varieties. Notably, the cowpea variety Lvlinghongshuai developed the most significant HR compared to the other tested varieties (Supplementary Fig. 1a). As we don’t know which viral protein of SMV can be recognized as Avr by NLRs from *Rsv1* locus, we constructed the full-length infectious clone of SMV using the SMV-SC7 isolate available in our laboratory to express all viral genes. To better visualize the viral infection, we also inserted a green fluorescent protein (GFP) reporter gene into the genome sequence of SMV. We then introduced the infectious clone of SMV-GFP in the cowpea cotyledons via an *Agrobacterium*-mediated transient expression. The results showed that the infection of SMV-GFP was successfully detected in all six cowpea varieties through both fluorescence imaging and immunoblotting analysis (Fig. 1b, c). Similar to the results of CMV 2a expression, the cowpea variety Lvlinghongshuai exhibited significantly higher SMV-GFP protein accumulation levels in cotyledon compared to the other five varieties (Fig. 1c). This cowpea variety was therefore used in subsequent studies.

**Fig. 1.**
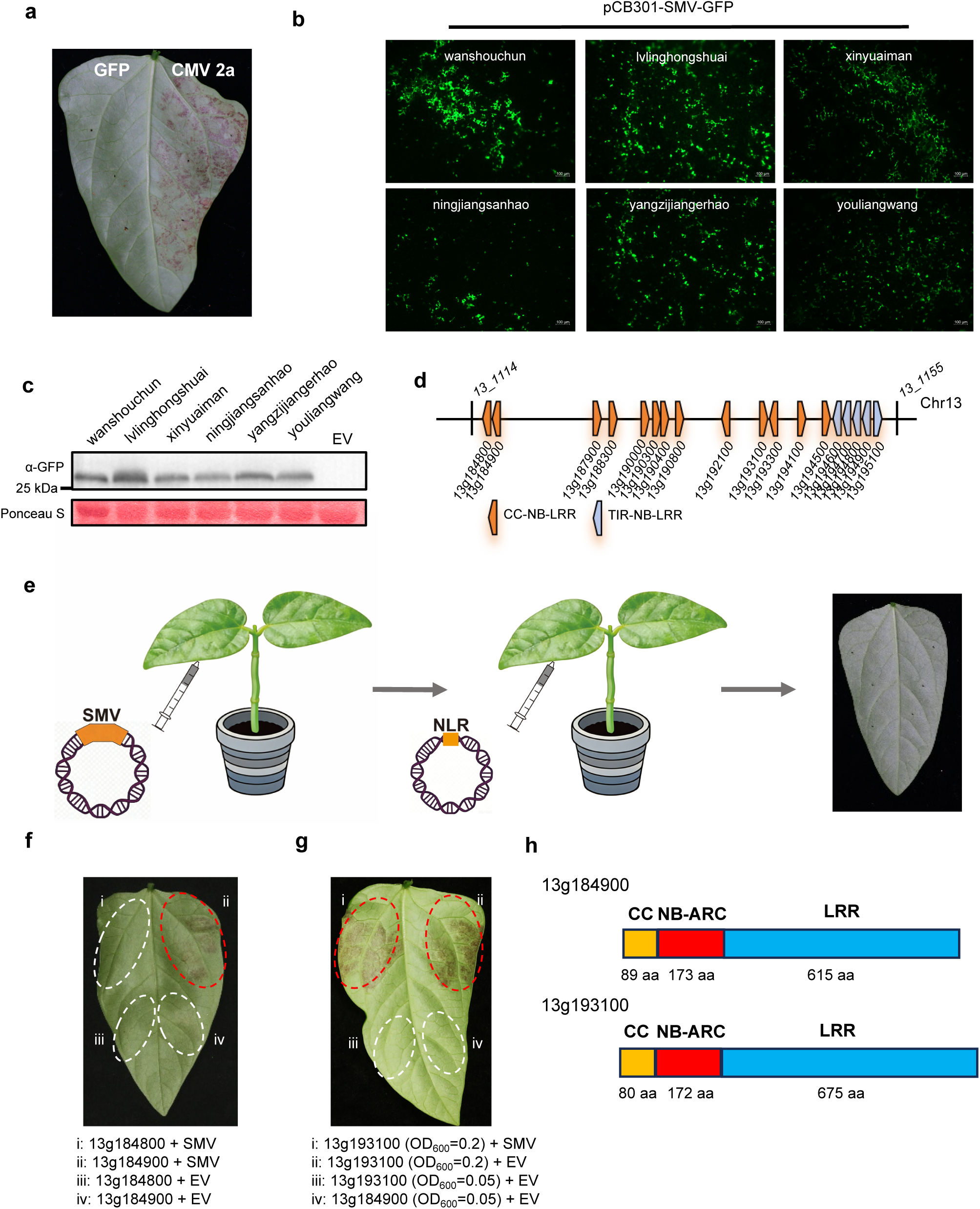
Establishment of a rapid and efficient transient expression system for screening of NLR genes from the soybean *Rsv1* locus. a. HR phenotype in cowpea cotyledons transiently expressing the CMV Avr protein 2a via *Agrobacterium* infiltration. pCambia2300 empty vector (EV) served as the negative control. b. Fluorescence images showing SMV-GFP infection in cotyledon tissues of six cowpea varieties after agro-infiltration with the full-length infectious clone of SMV carrying GFP reporter. Images were captured using an inverted fluorescence microscope. Cowpea cotyledons were agroinfiltrated with the SMV-GFP infectious clone and fluorescence imaged at 24 h post-infiltration. c. Immunoblot detection of GFP expressed from SMV in cotyledons of six cowpea varieties following infiltration with the SMV-GFP infectious clone. EV-infiltrated samples were used as negative controls. Ponceau S staining served as the loading control. Cowpea cotyledons were agroinfiltrated with the SMV-GFP infectious clone and assayed at 24 h post-infiltration. d. Schematic representation of the *Rsv1* locus showing the arrangement of 18 NLR genes in soybean cv Suweon 97. CC-NB-LRR-type and TIR-NB-LRR-type NLR genes are depicted in brown and grey colours, respectively. The locus is located on chromosome 13. e. Schematic overview of the screening strategy used to identify functional NLRs that recognize SMV. Cowpea cotyledons are infiltrated first with *Agrobacterium* carrying the SMV-GFP infectious clone, followed at 48 hpi by a second infiltration with *Agrobacterium* carrying individual NLR constructs. f. HR phenotypes in cowpea cotyledons co-expressing the NLR gene *13g184800* or *13g184900* with SMV-GFP at 5 days post infiltration. g. HR phenotypes of *13g193100* co-expressed with or without SMV-GFP in cowpea leaves at 5 dpi. h. Domain organization of NLR proteins encoded by *13g184900* and *13g193100*.

Analysis of genetic mapping and the soybean reference genome indicates that 18 clustered NLR genes are predicted within the *Rsv1* locus, including *13g184800*, *13g184900*, *13g187900*, *13g188300*, *13g190000*, *13g190300*, *13g190400*, *13g190800*, *13g192100*, *13g193100*, *13g193300*, *13g194100*, *13g194500*, *13g194600*, *13g194700*, *13g194800*, *13g194900* and *13g195100* (Fig. 1d). We individually screened these 18 NLR genes from *Rsv1* locus of Suweon 97 to determine whether they recognize SMV and induced HR response through the *Agrobacterium*-mediated transient expression in cowpea (Fig. 1e). In this assay, recognition of SMV by a transiently expressed functional NLR is expected to trigger HR cell death in cowpea leaf, allowing rapid identification of functional NLR genes without the need of stable soybean transformation. Following screening all 18 NLRs using this assay system, we found that the cowpea leaves co-expressing the NLR gene *13g184900* and SMV-GFP developed a clear HR cell death (Fig. 1f). Another NLR gene *13g193100* also induces HR with co-expression of SMV-GFP; however, it also induces HR in the absence of SMV-GFP, suggesting strong autoimmunity for this NLR (Fig. 1g). In contrast, none of the other NLR genes triggered detectable HR (Supplementary Fig. 1b). Both of *13g184900* and *13g193100* encode coiled-coil NLRs (CNL) resistance proteins, consisting of 1130 and 1189 amino acid residues, respectively (Fig. 1h). The two genes exhibit 73.2% and 68.9% sequence identity at the nucleotide and amino acid levels, respectively. Collectively, we established an HR-based rapid screening system for identification of NLR genes within complicated NLR locus and demonstrate that this approach can efficiently pinpoint the specific NLR responsible for SMV recognition among multiple NLR candidates in the *Rsv1* locus.

### NLR encoded by *13g184900* mediates the effective resistance to SMV

Next, we employed the cowpea transient expression system to examine whether the NLR gene *13g184900* or *13g193100* confers the resistance against SMV. The infectious clones of SMV-GFP and potato virus X (PVX)-GFP was co-expressed with the construct p2300-13g184900 under 35S promoter or pCambia empty vector (EV) control in cowpea cotyledon leaves. We anticipated that if NLR encoded by *13g184900* specifically recognizes SMV, it would confer the resistance to SMV but not PVX. Both fluorescence imaging and immunoblotting results showed that co-expression of *13g184900* significantly reduced the distribution and intensity of GFP expressed from SMV-GFP in cowpea leaves compared to co-expression of the EV control (Fig. 2a, b). In contrast, 13g184900 did not inhibit GFP expression from PVX-GFP (Fig. 2c, d). These results suggest that 13g184900 specifically mediates resistance against SMV, but not PVX infection. Similarly, we also tested the antiviral effect of *13g193100.* However, high-level overexpression of *13g193100* triggered strong autoimmunity, which confers the non-specific resistance. When we reduced the OD_600_ of the *Agrobacterium* carrying *13g193100* to 0.05, no significant antiviral effect on SMV-GFP infection was observed (Supplementary Fig. 2).

**Fig. 2.**
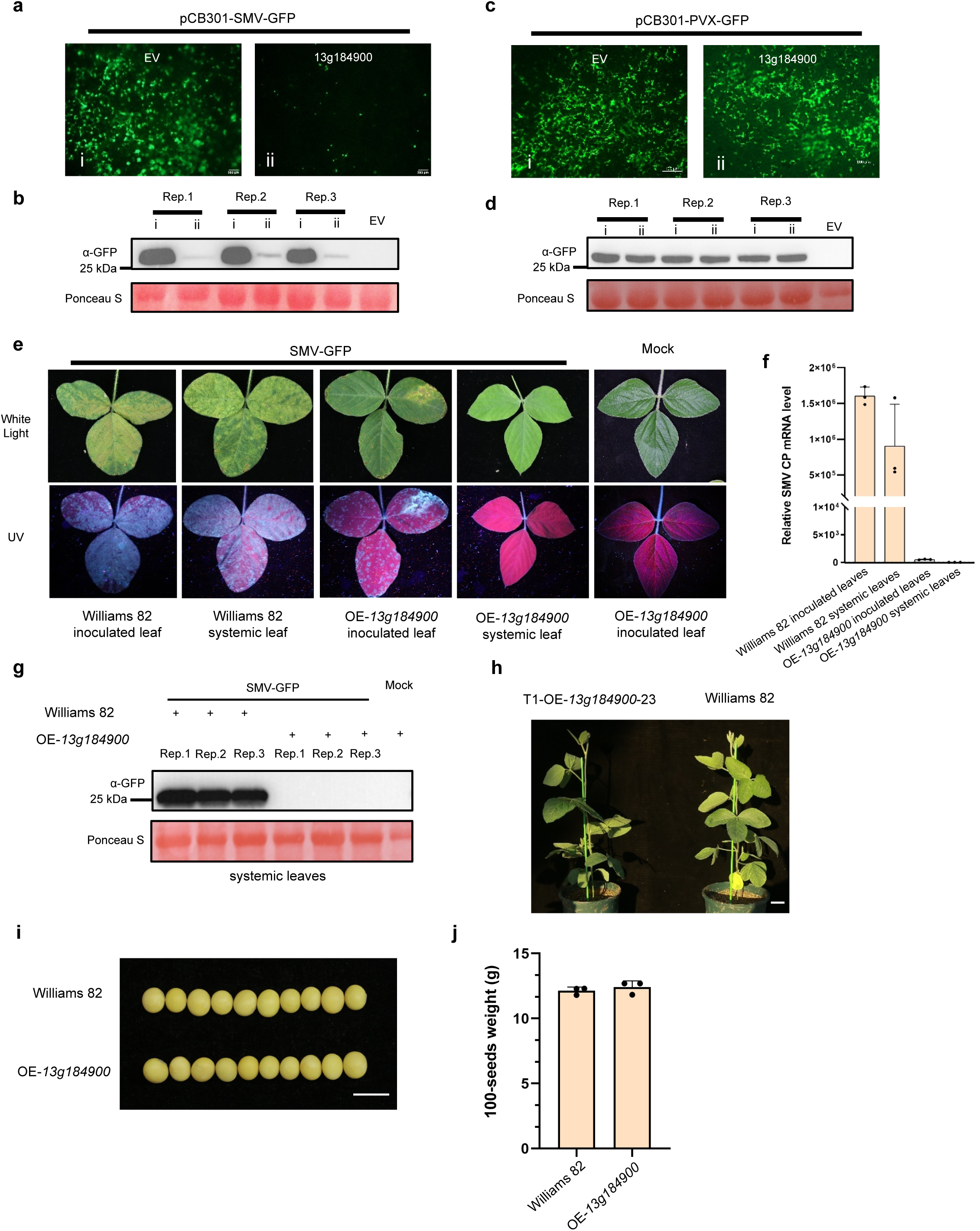
*13g184900* confers the effective resistance to SMV. a, b. Fluorescence imaging (a) and immunoblotting (b) of cowpea cotyledons co-expressing SMV-GFP infectious clone with 13g184900 or empty vector (EV) control. Cowpea cotyledons were fluorescence imaged at 48 h post-infiltration. Rep. 1–3 indicate three independent biological replicates. Ponceau S staining was used as a loading control. c, d. Fluorescence imaging (a) and immunoblotting (b) of cowpea cotyledons co-expressing PVX-GFP with 13g184900 or EV control. Ponceau S staining was used as a loading control. e. Inoculated leaves and systemic leaves symptoms in William 82 and OE-*13g184900* inoculated by SMV-GFP under white light (upper panel) and ultraviolet (UV) light (lower panel). OE-*13g184900* plants inoculated with PB were served as the mock control. f. Relative expression of SMV CP mRNA in local and systemic leaves of SMV-inoculated William 82 and OE-*13g184900* plant leaves determined by RT-qPCR. Data are presented as means ± SD, *n* = 3 independent biological replicates. g. Immunoblot analysis of SMV-GFP accumulation in systemic leaves of William 82 and OE-*13g184900* plants at 14 dpi. Ponceau S staining was served as a loading control. h. Phenotype of 6-week-old transgenic line (OE-*13g184900*-23) and the non-transgenic soybean cultivar Williams 82. Scale bar: 5cm. i. Seed morphology of OE-*13g184900* plants and Williams 82. Scale bar: 1cm j. 100-seed weight of OE-*13g184900* plants and Williams 82.

To further evaluate whether the *13g184900* confers SMV resistance in soybean, a binary expression vector containing this NLR gene driven by its native promoter was constructed and transformed into the SMV-susceptible cultivar Williams 82^39^. Twelve T_0_ transgenic plants were generated, and four T₁ transgenic lines were obtained, with line T₁-23 showing the highest expression level of *13g184900*. We also transformed *13g193100* into soybean. In contrast, transgenic plants expressing *13g193100* consistently developed chlorosis and necrosis during the differentiation stage, and failed to regenerate into plants. Consequently, no soybean transgenic lines expressing *13g193100* were successfully generated. Therefore, we utilized the soybean transgene lines harboring the *13g184900* (OE-*13g184900*) in the subsequent experiments.

Three-week-old Williams 82 and OE-*13g184900* were mechanically inoculated with extracts from SMV-GFP infected leaves. At 10 days post-inoculation (dpi), typical yellow-green mosaic symptoms were appeared on inoculated leaves of Williams 82, whereas OE-*13g184900* inoculated leaves exhibited typical HR cell death induced by SMV infection (Fig. 2e). By 17 dpi, mosaic symptoms also appeared on the upper new leaves of Williams 82, indicating that SMV-GFP had established systemic infection. In contrast, no mosaic-related symptoms were detected on the newly developed leaves of the OE-*13g184900* (Fig. 2e). Under ultraviolet (UV) light, green fluorescence was detected in both inoculated and systemic leaves of Williams 82, whereas no such fluorescence was detectable in the leaves of OE-*13g184900* (Fig. 2e). Viral accumulation assays showed that neither coat protein (CP) nor GFP of SMV-GFP was detected in systemic leaves of the transgenic plants (Fig. 2f, g). These results suggest that *13g184900* can effectively confer SMV resistance in soybean.

We also investigated whether the introduction of *13g184900* cause any fitness cost to soybean. Under identical growth conditions, six-week-old OE-*13g184900* showed no significant difference in plant height or growth fitness cost compared with the non-transgenic Williams 82 (Fig. 2h). In addition, the OE-*13g184900* showed no significant difference in seed morphology and seed size compared with the non-transgenic plants, and no significant difference was detected in 100-seed weight (Fig. 2i, j).

### NLR encoded by *13g184900* recognizes P3 protein from all SMV G1-G7 strains as Avr protein

To determine the Avr protein recognized by NLR encoded by *13g184900*, we individually cloned all 11 protein-encoding genes of SMV-SC7 and co-expressed each with *13g184900*. Co-expression of 13g184900 with the SMV P3 protein triggered strong HR, whereas no HR was detected in co-expression with other SMV genes, indicating that SMV P3 is the Avr protein recognized by the NLR encoded by *13g184900* (Fig. 3a and Supplementary Fig. 3). To further dissect the molecular mechanism underlying their immune recognition, we performed a panel of *in vivo* and *in vitro* interaction assays to verify the interaction between 13g184900 and P3. Co-immunoprecipitation (Co-IP) assays confirmed the interaction between the full-length 13g184900 NLR protein and SMV P3 *in vivo* (Fig. 3b). The yeast two-hybrid (Y2H) assays and firefly luciferase complementation imaging (LCI) assays further showed that the coiled-coil (CC) domain of this NLR directly interacted with the P3 protein (Fig. 3c, d). Furthermore, no interaction was detected between the NB-ARC or LRR domain of 13g184900 and the P3 protein (Supplementary Fig. 4a-c).

**Fig. 3.**
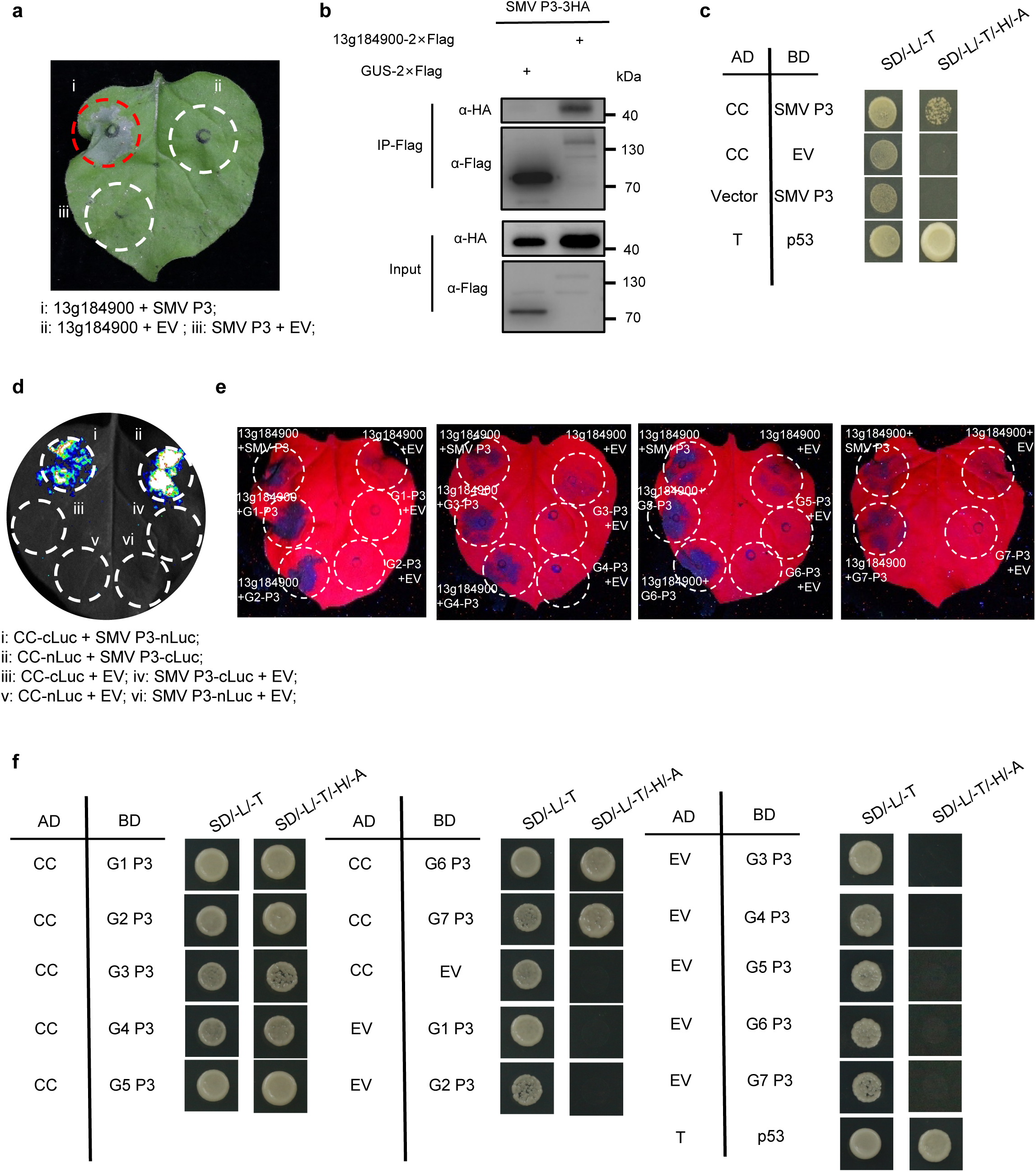
The NLR encoded by *13g184900* recognizes the P3 protein from all SMV G1–G7 strains as the Avr protein. a. HR phenotypes in *N. benthamiana* leaves transiently co-expressing *13g184900* and SMV P3. The P3 gene was cloned from the laboratory-maintained SMV-SC7 isolate. b. Co-immunoprecipitation (Co-IP) assays showing the interaction between 13g184900-2×Flag and SMV P3-HA co-expressed in *N. benthamiana* . GUS-2×Flag was included as a negative control. Proteins were immunoprecipitated with anti-Flag beads, and immunoblots were probed with anti-HA and anti-Flag antibodies, respectively. c. Yeast two-hybrid (Y2H) assays showing the interaction between the CC domain of 13g184900 and SMV P3. Empty vector (EV) and SV40 large T-antigen (T)/p53 served as negative and positive controls, respectively. d. Firefly luciferase complementation (LCI) assays showing the interaction between the CC domain of 13g184900 and SMV P3 in *N. benthamiana* leaves. e. HR phenotypes in *N. benthamiana* leaves co-expressing 13g184900 with P3 proteins from SMV G1 to G7 strains, respectively. These P3 genes from SMV G1 to G7 strains was chemically synthesized. These *N. benthamiana* leaves was photoed by UV light. f. Y2H assays showing that the CC domain of 13g184900 interacts with P3 proteins from all seven SMV strains (G1–G7). Empty vector (EV) and SV40 large T-antigen (T)/p53 served as negative and positive controls, respectively.

*Rsv1* locus of Suweon 97 can confer broad-spectrum resistance to SMV G1-G7 strains^35,40^. We chemically synthesized the P3 genes from SMV G1 to G7 strains^41,42^ and co-expressed each with *13g184900*, to examine whether *13g184900* is the NLR gene of *Rsv1* locus in Suweon 97 providing the broad-spectrum resistance to all SMV G1-G7 strains. Co-expression of 13g184900 with P3 from each strain all elicited HR (Fig. 3e), indicating that *13g184900* is the NLR gene that confers the broad-spectrum resistance to SMV in soybean. Furthermore, we demonstrated that CC domain of 13g184900 interacts with the P3 protein of all seven SMV strains (Fig. 3f). The nucleotide sequences of the P3 proteins from the seven SMV strains are highly conserved (≥88.89% identity) (Supplementary Fig. 5).

### NLR encoded by *13g184900* also mediates resistance to another potyvirus BCMV via recognizing viral P3 protein

Previous studies reported that the Suweon 97 confers resistance not only to SMV but also to BCMV^43^. To determine whether *13g184900* also confers resistance to BCMV, we constructed a BCMV infectious clone carrying a GFP gene (BCMV-GFP). We expressed the BCMV-GFP in cotyledons of the cowpea cultivar via the *Agrobacterium*-mediated transient system. The result showed that this infectious clone can successfully express in cowpea (Supplementary Fig. 6). We then co-expressed 13g184900 with BCMV-GFP infectious clone in cowpea cotyledons. The results showed that co-expression of 13g184900 and BCMV-GFP triggered HR cell death in cowpea leaves and strongly suppressed BCMV-GFP accumulation, indicating that *13g184900* effectively recognizes and confers the resistance against BCMV (Fig. 4a-c). We then inoculated the OE-*13g184900* and Williams 82 with BCMV-GFP. Inoculated leaves of transgenic plants developed localized necrotic lesions, while newly developed systemic leaves remained symptom-free (Fig. 4e). In contrast, non-transgenic plants exhibited typical mosaic symptoms on systemic infected leaves (Fig. 4e). Consistent with the phenotype, BCMV-GFP was not detected in upper new leaves of OE-*13g184900* by Western blot analysis (Fig. 4f).

**Fig. 4.**
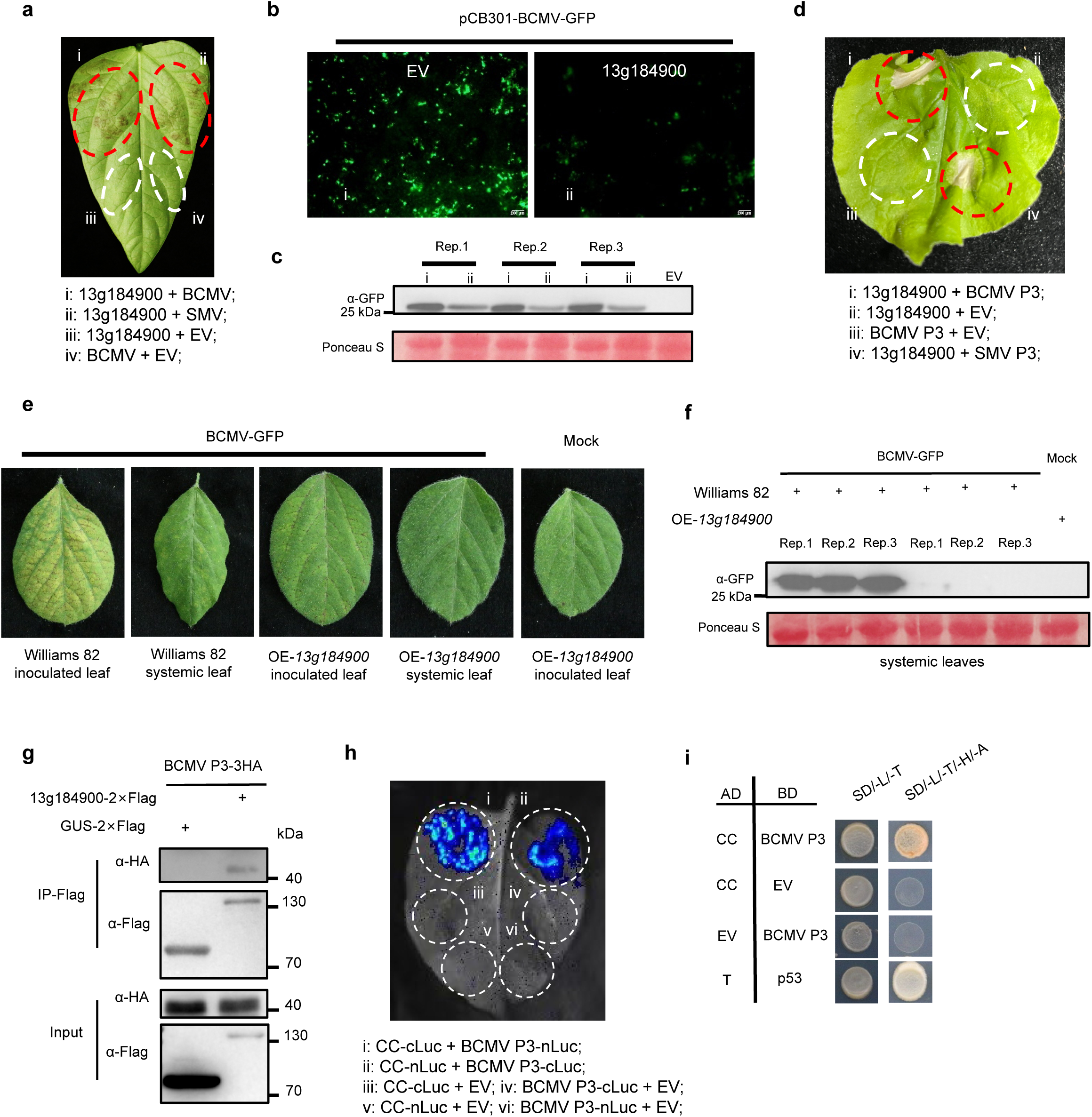
NLR encoded by *13g184900* also mediates resistance to another potyvirus BCMV via recognizing viral P3 protein. a. HR phenotypes in cowpea leaves transiently co-expressing 13g184900 with BCMV-GFP infectious clone. b, c. Fluorescence imaging (b) and immunoblotting (c) analysis of BCMV-GFP accumulation in cowpea cotyledons co-expressing BCMV-GFP infectious clone with 13g184900 or empty vector (EV) control. Cowpea cotyledons were fluorescence imaged at 48 h post-infiltration. Rep. 1–3 indicate three independent biological replicates. Ponceau S staining was used as a loading control. d. HR phenotypes in *N. benthamiana* leaves co-expressing 13g184900 with BCMV P3. e. Phenotypes of local leaf and systemic leaf of Williams 82 and OE-13g184900 plants inoculated with BCMV-GFP. Plants inoculated with PB were served as the mock control. f. Immunoblot analysis of BCMV-GFP accumulation in systemic leaves of Williams 82 and OE-*13g184900* plants at 14 dpi. Rep. 1–3 represent three independent biological replicates. Ponceau S staining was used as the loading control. g Co-IP assay showing the interaction between full-length 13g184900-2×Flag and BCMV P3-3HA co-expressed in *N. benthamiana*. GUS-2×Flag was included as a negative control. Proteins were immunoprecipitated with anti-Flag beads, and immunoblots were probed with anti-HA and anti-Flag antibodies, respectively. i. LCI assay showing the interaction between the CC domain of 13g184900 and BCMV P3. j. Y2H assay demonstrating the direct interaction between the CC domain of 13g184900 and BCMV P3. Empty vector (EV) and SV40 large T-antigen (T)/p53 served as negative and positive controls, respectively.

Furthermore, we cloned the gene encoding BCMV P3 protein and found that 13g184900 can also recognize BCMV P3 and induces HR (Fig. 4d). Co-IP assays confirmed the interaction between the full-length 13g184900 NLR protein and BCMV P3 *in vivo* (Fig. 4g). LCI assays and Y2H showed that the CC domain of the *13g184900*-encoded NLR protein interacted with the BCMV-P3 protein (Fig. 4h, i). Sequence alignment analysis revealed that BCMV P3 shares ∼58.7% nucleotide sequence identity with SMV P3 (Supplementary Fig. 7). Together, these results demonstrate that *13g184900* encodes a NLR that recognizes P3 from all G1–G7 SMV strains and also confers the resistance against the related potyvirus BCMV by interacting with viral P3 protein.

### Evolutionary origin of the NLR gene *13g184900*

To explore the evolutionary origin and functional conservation of *13g184900* in Suweon 97, we constructed two phylogenetic trees based on its homologs from wild soybean (*Glycine soja*) and cultivated soybean. As shown in Fig. 5a, *13g184900* clustered tightly with homologs from wild soybeans YSD56, PI483463, and Yongin-0 ( ≥ 97.84% identity). Because NLR protein encoded by *13g184900* recognizes SMV/BCMV Avr proteins via its CC domain, we also performed phylogenetic analysis using the CC domain-encoding sequences of homologous genes from Suweon 97 and diverse wild soybean accessions (Fig. 5b). The result exhibited an overall evolutionary pattern highly consistent with that constructed from full-length coding sequences. Among them, *13g184900* from Suweon 97 clustered tightly with its orthologous genes on Chr 11 and Chr 13 of wild soybean into a clade, with sequence identities ranging from 95.3% to 99.67%. The highest sequence identity (99.67%) was observed between *13g184900* and its ortholog on Chr 11 of wild soybean species YSD56 originated from China (Fig. 5a).

**Fig. 5.**
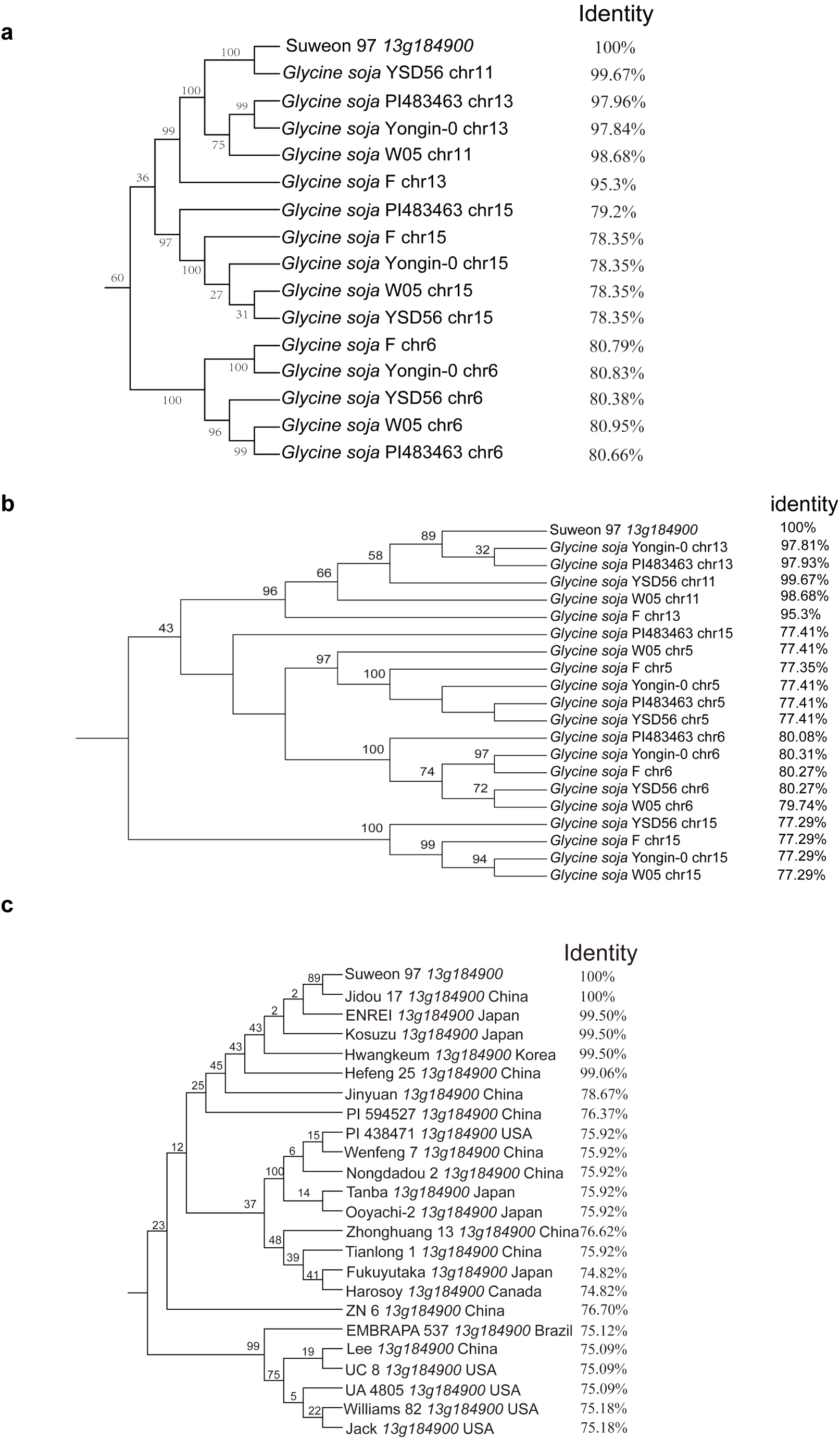
Phylogenetic analyses of the evolutionary origin and diversification of the NLR gene *13g184900* in soybean. a. Phylogenetic tree of full-length *13g184900* homologs from wild soybean (*Glycine soja*) accessions. Identity values between each pair of sequences are shown on the right. b. Phylogenetic tree of full-length *13g184900* homologs from cultivated soybean (*Glycine max*) worldwide. Identity values between each pair of sequences are shown on the right. c. Phylogenetic tree built using CC-domain coding sequences of *13g184900* homologs from wild soybean accessions. Bootstrap support values are labelled at each internal node of all trees.

In cultivated soybeans, *13g184900* clustered with homologs from Jidou 17, ENREI, and Kosuzu (>99.5% identity), while other cultivars like Williams 82 and Jack formed a clade (75.09%–78.67% identity) (Fig. 5c). This gene exhibits exceptionally high sequence similarity across its ortholog in diverse soybean cultivars, indicating strong evolutionary conservation that is critical for its disease resistance.

## Discussion

Soybean is one of the most economically important crop worldwide. Among soybean pathogens, SMV represents one of the most impactful and widely distributed viral agents. Accordingly, the cloning and functional characterization of SMV resistance genes are critically important for soybean disease resistance breeding. In this study, we established a rapid screening system for the identification of NLR genes from soybean complex NLR locus based on transient agro-infiltration of cowpea cotyledons. Using this system, we identified a broad-spectrum NLR resistance gene, *13g184900*, from the highly complicated *Rsv1* locus, recognizing all SMV G1–G7 strains as well as another potyvirus BCMV. Stable expression of *13g184900* in soybean conferred resistance to both SMV and BCMV.

Our cowpea cotyledon-based screening system allows rapid functional screening of the 18 NLR genes at the *Rsv1* locus from Suweon 97 via HR-induced cell death, and the intensity and spatial distribution of GFP fluorescence. This cowpea-based platform is also well suited for screening NLRs and AVRs that engage in indirect interactions, because the legume host factors required for such indirect recognition would be missed in the *N. benthamiana*-based screening system^44–46^. Beyond its utility in antiviral NLR gene discovery, this screening system can be extended to mine NLR genes against other soybean pathogens, including *Phytophthora sojae*, rust fungi, and nematodes, thereby providing an efficient technical platform for the functional dissection of complex NLR locus in soybean. Moreover, this system can be applied to screen NLR genes from other leguminous crops, such as common bean, faba bean, cowpea, snow pea, and mung bean. Additionally, since SMV-GFP and BCMV-GFP can accumulate to high levels in cowpea cotyledons, the transient expression system established in this study can also be applied to investigate the replication, movement, and host resistance mechanisms of SMV and BCMV.

We also identified that the NLR gene *13g184900* recognizes SMV P3 as its Avr protein. However, the other NLR gene identified, *13g193100*, exhibits strong autoimmunity, thus preventing identification of its Avr determinant. Previous studies showed that coordinated mutations in the P3 and HC-Pro proteins enable SMV to evade resistance mediated by the *Rsv1* allele from soybean cultivar PI96983, leading to the result that both P3 and HC-Pro act as Avr proteins recognized by this *Rsv1*^47–49^. The NLR gene *13g193100* may be responsible for HC-Pro recognition. We also found that NLR encoded by *13g184900* is capable of recognizing P3 from all seven SMV strains. Collectively, these results indicate that *13g184900* is the key resistance gene underlying broad-spectrum SMV resistance at the *Rsv1* in Suweon 97. However, as *Rsv1* alleles vary widely in genetic composition and genomic organization across different soybean cultivars, individual alleles may confer recognition of distinct Avr protein. The Avr determinants recognized by the *Rsv1* alleles in Suweon 97 and other cultivars need further characterizations.

SMV resistance is an essential trait of soybean resistance breeding in China. And SMV strains are diverse and highly variable, and single resistance genes frequently suffer breakdown^50^. To date, only two SMV resistance genes *Rsv4* and *Rsc4-3* have been cloned and functionally validated. These two genes fall into distinct resistance gene families, with pronounced differences in their antiviral mechanisms. *Rsv4* is not a canonical NLR-type resistance gene^51^. Its resistance is marked by delayed symptom development and restricted systemic viral spread rather than complete immunity, and can be overcome by specific viral mutants. *Rsc4-3* belongs to the CNL-type resistance gene and is allelic to the *Rsv3* locus candidate *NBS_C*, with partially overlapping genomic mapping intervals and conserved antiviral function against SMV. *Rsc4-3* can confer resistance to several SMV strains, but fails to confer resistance to strain SC15^52,53^. Therefore, the broad-spectrum SMV-resistant NLR gene *13g184900* identified in this study provides a valuable gene resource for breeding soybean varieties against different strains of SMV.

BCMV poses threats to other beans, too^20^. Hence, the NLR gene *13g184900* identified in this study can also be utilized in beans and other legumes to achieve resistance against BCMV. Whether *13g184900* confers broad-spectrum resistance to other potyviruses remains to be validated in future studies. Even if this NLR gene fails to recognize and resist other potyviruses, multiple studies have documented that the pathogen recognition range of NLR immune receptors can be expanded via genetic engineering or targeted mutagenesis^54–57^. The conserved Avr protein recognition mechanism of NLR encoded by *13g184900* described here provides an important basis and great potential for further utilization and engineering of this NLR gene to confer broad-spectrum resistance against multiple potyviruses in different crops.

Further sequence alignment analyses revealed that *13g184900* from soybean cultivar Suweon 97 shares 99.67% sequence identity with its homologs located on chromosome 11 of wild soybean accession YSD56. This indicated that this NLR gene originated from wild soybean originated in East China. The occurrence of these homologous genes on distinct chromosomes reflects the homeologous relationship between soybean chromosome 11 and 13, which arose from a whole-genome duplication (WGD) event^58–60^. Specifically, the NLR resistance gene cluster harboring the *Rsv1* locus on chromosome 13 has a corresponding homeologous NLR cluster in the syntenic region of chromosome 11. In terms of germplasm distribution, this gene is present in several soybean cultivars from East Asia but is absent in cultivars from other major soybean-producing regions included in this analysis. Our study identified the other NLR gene at the locus, *13g193100*, which triggers strong autoimmunity. Consequently, we were unable to generate stable transgenic soybean lines expressing this gene. Based on these results, we speculate that the fitness cost conferred by other NLR member like *13g193100* within the *Rsv1* locus may have prevented the introduction of *Rsv1* locus into cultivated soybeans in other major production regions during breeding history. However, there is no study systematically compared growth traits between soybean lines with and without the *Rsv1* locus. The evolutionary and breeding mechanisms underlying this distinct distribution pattern remain largely unexplored.

In summary, we developed an efficient system for screening 18 NLRs in highly complicated *Rsv1* locus in the soybean and identified *13g184900* as a broad-spectrum resistance gene recognizing all SMV G1-G7 strains and another potyvirus BCMV. Our findings further highlight the promising application value of *13g184900* for molecular resistance breeding across diverse soybean cultivars. And these findings lay an important and solid foundation for further NLR gene identification, breeding of broad-spectrum antiviral cultivars, and studies of NLR-AVR interactions in soybean.

## Methods

### Plant material and cultivation conditions

For the phenotyping assays, the seeds of Cowpea (including 6 commercial cowpea varieties: Lvlinghongshuai, Wanshouchun, Xinyuaiman, Ningjiangsanhao, Yangzijiangerhao, Youliangwang), *N.benthamiana*, and soybean (Williams 82 and *13g184900*-overexpression plants) were sown in a 1:1 mixture of Pindstrup Substrate (0-6 mm, pH = 6.0) and vermiculite, and cultivated in several environment-controlled greenhouse under a 16 h light (25 °C)/8 h dark (23 °C) photoperiod. To obtain overexpression lines, seedlings of soybean cultivar Williams 82 plants were grown in tissue culture medium in the environment-controlled chamber. The growth chambers were operated under a 14 h light / 10 h dark photoperiod, with an illumination intensity of 20,000 lux.

### Plasmid construction and soybean transformation

The full-length RNA of SMV strain SC7 was extracted from systemically infected leaves of soybean cultivar Williams 82 and reverse-transcribed into first-strand cDNA. The eGFP coding sequence was inserted between the NIb and CP coding regions of the SMV. The full-length SMV-GFP recombinant fragment was cloned into the binary vector pCB301 downstream of the CaMV 35S promoter using the ClonExpress II One Step Cloning Kit (Vazyme Biotech, Nanjing, China). The final infectious clone construct transformed into *Agrobacterium tumefaciens* strain GV3101 for subsequent viral inoculation assays. The full-length coding sequences of the individual NLR genes from the *Rsv1* locus and the viral protein of SMV and BCMV were amplified by RT-PCR, fused with HA or FLAG tags at their N-terminus or C-terminus, and cloned individually into pCambia2300S under the control of the 35S promoter. The full-length coding sequences of NLR genes *13g184900* and *13g193100* were inserted into the plant binary vector pFGC5941, under their native promoter, with the bar gene conferring glufosinate resistance as the selectable marker. These two overexpression constructs were introduced into *A. tumefaciens* strain AGL1 and used to generate transgenic soybean plants in the Williams 82 background.

Soybean genetic transformation was performed using an established cotyledonary-node transformation protocol with minor modifications^61,62^. Well-rooted T_0_ plants transplanted into pots containing a 1:1 (v/v) mixture of substrate and vermiculite, and grown in a greenhouse under the same photoperiod and temperature conditions. Then T_0_ plants were screened by herbicide leaf painting with 100 mg/l of glufosinate onto trifoliate leaves. Genomic DNA was extracted from glufosinate-tolerant plants, and positive transgenic lines were further confirmed by PCR amplification of the target NLR gene and the bar selection marker.

### Transient expression in cowpea and *N. benthamiana*

Fourteen-day-old seedlings with fully expanded cotyledons were used for agroinfiltration. *Agrobacterium tumefaciens* strain GV3101 carrying the target constructs (including the SMV-GFP, BCMV-GFP, PVX-GFP infectious clone, and NLR overexpression vectors) were cultured overnight in LB medium supplemented with 100 mg/L kanamycin and 50 mg/L rifampicinat 28 °C with shaking at 220 rpm. GV3101 cells were harvested by centrifugation at 5000 rpm for 5 min, then resuspended in agroinfiltration buffer (10 mM MES pH 5.6, 10 mM MgCl₂, 150 mM acetosyringone). The suspensions were incubated at 28 °C in the dark for 2–3 h before infiltration. For the screening system in cowpea, agrobacterium cultures carrying infectious clones of SMV-GFP or BCMV-GFP were first adjusted to an OD₆₀₀ of 1.0 and infiltrated into cowpea cotyledons. The inoculated plants were then incubated at 25 °C for 24h, followed by a second infiltration with agrobacterium cultures harboring NLR gene constructs adjusted to an OD₆₀₀ of 0.5. Infiltrated cowpeas were returned to the greenhouse under the same conditions. HR cell death phenotypes were visually recorded at 5 days post-infiltration (dpi). GFP fluorescence signals were observed and imaged using a fluorescence microscope at 36 hpi. *Agrobacterium*-mediated transient expression in *N. benthamiana* leaves was performed following the method of reference^63^.

### SMV/BCMV mechanical inoculation

To better visualize viral infection, we performed mechanical inoculation using the SMV-SC7-GFP infectious clone. *N.benthamiana* plants were first agroinoculated with the SMV-SC7-GFP infectious clone. Once systemic disease symptoms developed at 14 days post-inoculation, infected leaf tissues were harvested and ground in 1× phosphate buffer (137 mM NaCl, 2.7 mM KCl, 10 mM Na_2_HPO_4_, and 2 mM KH_2_PO_4_, pH 7.4). The crude homogenate was used immediately for inoculation to maintain viral infectivity. Soybean plants with fully expanded leaves (approximately two weeks after sowing) were used as inoculation materials. The surface of unifoliate leaves was lightly dusted with 600-mesh carborundum as an abrasive. The viral homogenate was gently rubbed across the leaf surface with a sterile gloved finger. After inoculation, residual inoculum and carborundum were rinsed off with sterile distilled water. Mock inoculation with phosphate buffer was set as the negative control. Mechanical inoculation of BCMV was performed following the same protocol described above for SMV.

### Co-IP assays

For Co- IP, *Agrobacterium tumefaciens* strains harbouring constructs encoding FLAG- tagged 13g184900 and HA-tagged SMV P3 (or the empty vectors as controls) were co- infiltrated into *N. benthamiana* leaves. At 24 hpi, 1 g of infiltrated leaf tissue was collected and ground in 2 mL of ice-cold extraction buffer (10% [v/v] glycerol, 25 mM Tris-HCl [pH 7.5], 1 mM EDTA, 150 mM NaCl, 10 mM dithiothreitol, 2% [w/v] polyvinylpolypyrrolidone, 0.5 % [v/v] Triton X-100, and complete protease inhibitor cocktail) using a mortar and pestle. The homogenate was centrifuged at 18,000 ×g for 30 min at 4 °C, and the supernatant was incubated with 30 μL of anti-FLAG agarose beads (Sigma-Aldrich, St. Louis, MO, USA) for 2 h at 4 ℃ with gentle rotation. The beads were then washed six times with IP buffer (50 mM Tris-Cl [pH 8.5], 100 mM NaCl, and 1 mM EDTA), resuspended in SDS-PAGE loading buffer (150 mM Tris-HCl [pH 6.8], 30 % glycerol, 6 % SDS, 0.3 % bromophenol blue, and 300 mM DTT), and boiled at 95 °C for 5 min. Eluted proteins were resolved by SDS-PAGE, transferred to PVDF membranes, and immunoblotted with anti-FLAG (Sigma-Aldrich, Cat. # A8592, 1:10,000) and anti-YFP (Sigma-Aldrich, Cat. # SAB4301138, 1:10,000) primary antibodies, followed by HRP-conjugated goat anti-rabbit secondary antibody (Sigma-Aldrich, Cat. # A0545, 1:10,000). Ponceau S staining of the membranes was used to confirm equal protein loading.

### RT–qPCR analysis

Total RNA was extracted from soybean leaves using a FastPure Universal Plant Total RNA Isolation Kit (Vazyme Biotech, Nanjing, China). The extracted RNA was reverse-transcribed into cDNA by HiScript III RT SuperMix (Vazyme Biotech, Nanjing, China).We performed qRT - PCR using ChamQ Universal SYBR qPCR Master Mix (Vazyme Biotech, Nanjing, China) on an ABI Prism 7500 Fast Real-Time PCR system (Life Technologies, Beverly, MA, USA). *GmActin11* were used as internal controls.

### Y2H assays

Y2H Gold yeast cells were co-transformed with different combinations of bait (pGBKT7/BD) and prey (pGADT7/AD) constructs. Transformants were first selected on synthetic defined medium lacking tryptophan and leucine (SD/-Trp-Leu) and incubated at 28 °C for 3 days. The yeast cells were diluted in ddH_2_O at ratios of 1:1 (v/v) and grown on SD/-Trp-Leu-His-Ade media for 3 days at 28 °C. Yeast transformed with BD-P53 and AD-SV40T was used as a positive control.

### LCI assay

The coding sequences of the CC, NB-ARC, and LRR domains of Glyma13g184900, as well as the full-length P3 coding sequences of SMV and BCMV, were individually cloned into the N-terminal firefly luciferase (nLUC) and C-terminal firefly luciferase (cLUC) vectors, respectively. All recombinant constructs were verified by Sanger sequencing and transformed into *A.tumefaciens* strain GV3101. For transient co-expression, equal volumes of the corresponding *A. tumefaciens* suspensions (OD₆₀₀ = 1.0) were mixed and infiltrated into leaves of 4–5-week-old *N. benthamiana* plants. At 24 hours post-infiltration, 150 μg/mL of luciferin potassium salt (Yeasen, Shanghai, China) was evenly sprayed onto the infiltrated leaf areas. After 5–10 min of dark incubation to enable substrate penetration and reduce background noise, luciferase luminescence signals were detected and imaged using a low-light cooled CCD imaging system (Tanon, Shanghai).

### Sequence alignment and phylogenetic tree construction

The full-length coding sequence of *13g184900* was cloned from soybean cultivar Suweon 97 in this study. Homologous NLR sequences from wild soybean accession (including YSD56, PI483463, Yongin-0, W05, F) and soybean cultivar (including ENREI, Kosuzu, Hwangkeum, Hefeng 25, Jinyuan, PI 594527, PI 438471, Wenfeng 7, Nongdadou 2, Tanba, Ooyachi-2, Zhonghuang 13, Tianlong 1, Fukuyutaka,, Harosoy, ZN 6, EMBRAPA 537, Lee, UC 8, UA 4805, Williams 82, Jack) were retrieved from the National Center for Biotechnology Information (NCBI) database and published wild soybean genome assemblies, respectively. Multiple sequence alignment was performed using MAFFT v7.0 with default parameters. Pairwise nucleotide sequence identity was calculated using the EMBOSS Needle tool based on the aligned sequences. The phylogenetic tree of *13g184900* and homologs was built using a Maximum Likelihood Estimate implanted and visualized in MEGA-X.

## Acknowledgments

Funding for this project was provided by the National Key Research and Development Program of China (2022YFD1401200 and 2023YFD1401000), the National Natural Science Foundation of China (32430088 and 32220103008), the Fundamental and Interdisciplinary Disciplines Breakthrough Plan of the Ministry of Education of China (JYB2025XDXM703), the Joint Research Program of State Key Laboratory of Agricultural and Forestry Biosecurity (No. SKLJRP2506), the Key Projects of YNTC (2025530000241001) and Yunnan Seed Laboratory (Grant No. 202205AR070001).

## Author contributions

T.X.R. and W.Y.C. supervised the work. T.X.R. and Z.H.Y. designed the study and wrote the manuscript. Z.H.Y. and G.B. performed the majority of the experiments and data analysis. L.J.Q. and Z.Y.Y. assisted in the generation of transgenic soybean lines. Z.Y.X., Y.T.Q., H.P.L., T.Y., W.M.H. contributed to vector construction and plant cultivation. G.L. helped with Sequence alignment and phylogenetic tree construction. C.X.Y. and C.X. provided the soybean seed materials used in this study. L.K. and Z.H.J. helped with pathogen inoculation assays. X.Y. contributed valuable advice on study design. D.K.X. provided technical expertise for soybean transformation.

## Supplementary Figure legends

**Supplementary Fig. 1.** Screening for NLR candidate genes from soybean *Rsv1* locus via transient agro-infiltration in cowpea cotyledons. a. HR phenotypes of six different cowpea varieties upon transient expression of CMV Avr 2a protein via agroinfiltration. pCambia2300 empty vector (EV) served as the negative control. b. HR phenotypes in cowpea cotyledons co-expressing the NLR gene *13g187900*, *13g188300*, *13g190000*, *13g190300*, *13g190400*, *13g190800*, *13g192100*, *13g193100*, *13g193300*, *13g194100*, *13g194500*, *13g194600*, *13g194700*, *13g194800*, *13g194900* and *13g195100* with SMV-GFP at 5 days post infiltration.

**Supplementary Fig. 2.** Low-level expression of *13g193100* shows no significant antiviral effect to SMV-GFP. The *13g193100*-carrying *Agrobacterium* suspension was adjusted to an OD₆₀₀ of 0.05 for infiltration. Fluorescence imaging of cowpea cotyledons co-expressing SMV-GFP infectious clone with 13g184900 or empty vector (EV) control. Rep. 1–3 indicate three independent biological replicates. Ponceau S staining was used as a loading control.

**Supplementary Fig. 3.** HR phenotypes in *N. benthamiana* leaves transiently co-expressing *13g184900* and other 10 SMV encoded proteins (including P1, HC-Pro, 6K1, CI, 6K2, VPg, NIa-Pro, NIb, CP and P3N-PIPO).

**Supplementary Fig. 4.** The NB-ARC and LRR domains of the NLR protein encoded by *13g184900* do not interact with the SMV P3 protein. a,b. Firefly luciferase complementation imaging (LCI) assays in *N. benthamiana* leaves. The NB-ARC (a) or LRR (b) domain of 13g184900 was fused to the N-terminal (nLuc) or C-terminal (cLuc) half of luciferase and co-expressed with SMV P3 fused to the complementary luciferase half or with an empty vector (EV) control, in the combinations indicated. c. Yeast two-hybrid (Y2H) assays shows that the NB-ARC domain or LRR domain of 13g184900 doesn’t directly interact with SMV P3. Empty vector (EV) and SV40 large T-antigen (T)/p53 served as negative and positive controls, respectively.

**Supplementary Fig. 5.** Phylogenetic analysis and sequence identity analysis of P3 coding sequences from seven SMV strains. The nucleotide sequences of P3 from SMV strains G2-G7 were compared with strain G1. Percent identity relative to G1 is shown for each strain. The nucleotide sequences encoding the P3 protein of SMV strains G1–G7 were downloaded from the National Center for Biotechnology Information (NCBI) database.

**Supplementary Fig. 6.** Expression of BCMV-GFP in cowpea cotyledons. a. Fluorescence images showing BCMV-GFP infection in cotyledon tissues of six cowpea varieties after agro-infiltration with the full-length infectious clone of BCMV carrying GFP reporter. Images were captured using an inverted fluorescence microscope. Cowpea cotyledons were agroinfiltrated with the SMV-GFP infectious clone and fluorescence imaged at 24 h post-infiltration. b. Immunoblot detection of GFP expressed from BCMV in cotyledons of six cowpea varieties following infiltration with the SMV-GFP infectious clone. EV-infiltrated samples were used as negative controls. Ponceau S staining served as the loading control.

**Supplementary Fig. 7.** Sequence alignment of BCMV P3 with SMV G1-G7 P3. The BCMV P3 coding sequence (1,041 bp) and the P3 sequences of seven SMV strains were aligned at nucleotide levels.

