## Supplementary Figures for "NLR from soybean *Rsv1* locus confers broad-spectrum resistance to soybean mosaic virus G1-G7 strains by recognizing viral P3 protein"

a

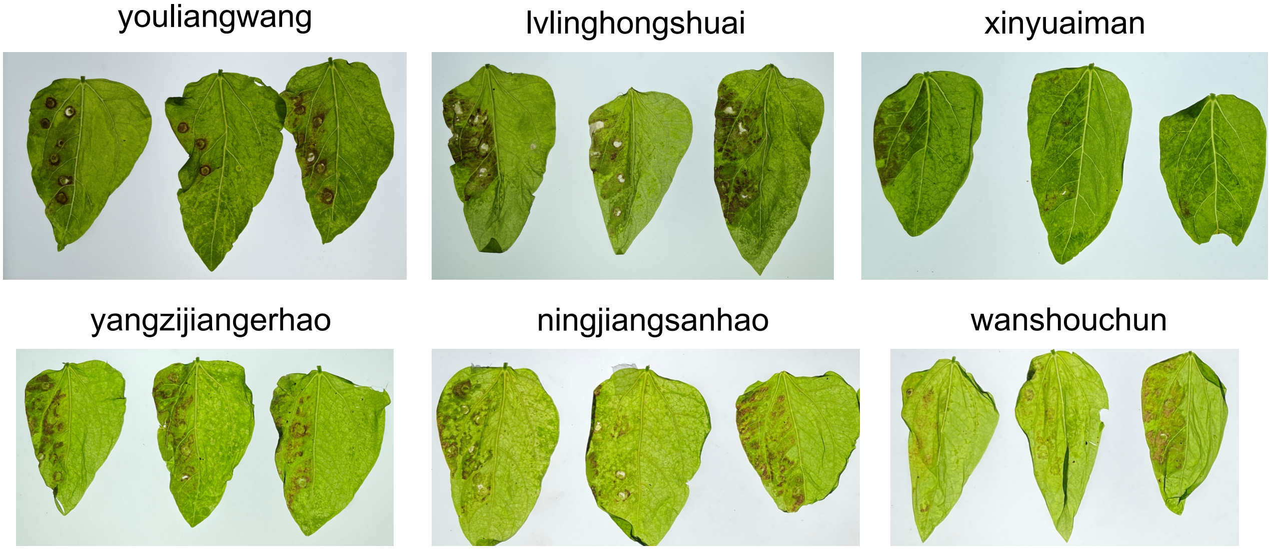


b

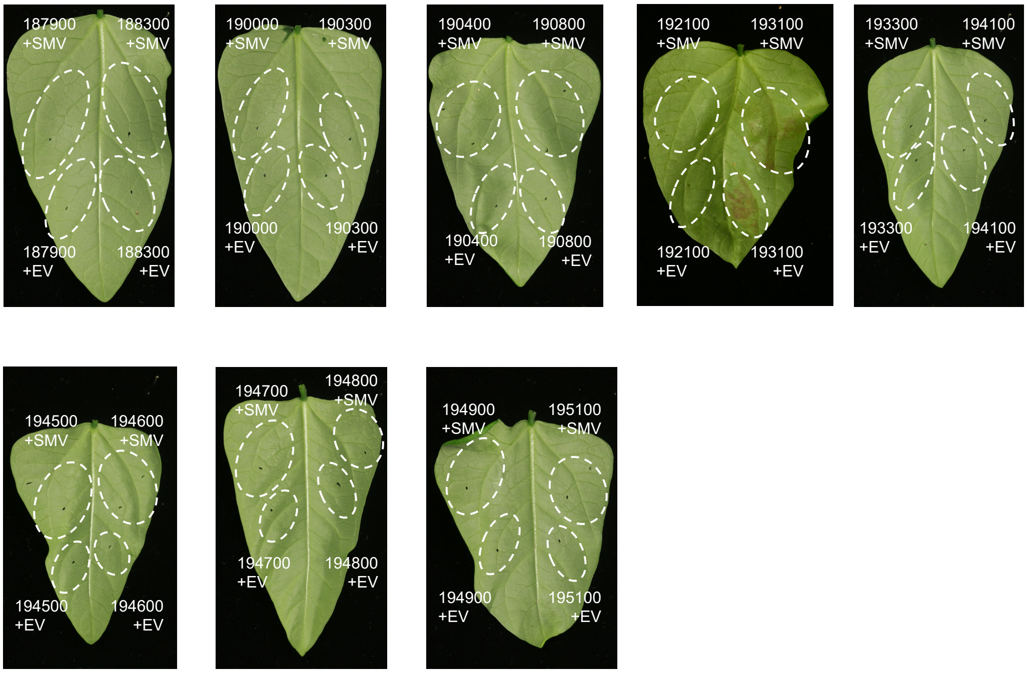


Supplementary Fig. 1

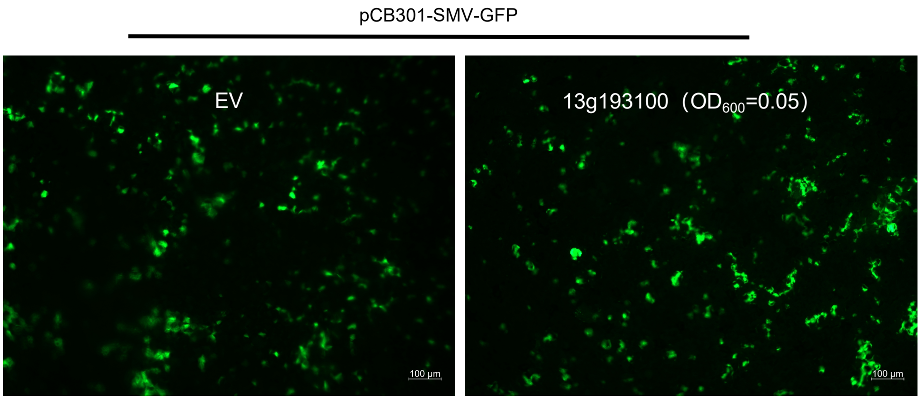


Supplementary Fig. 2

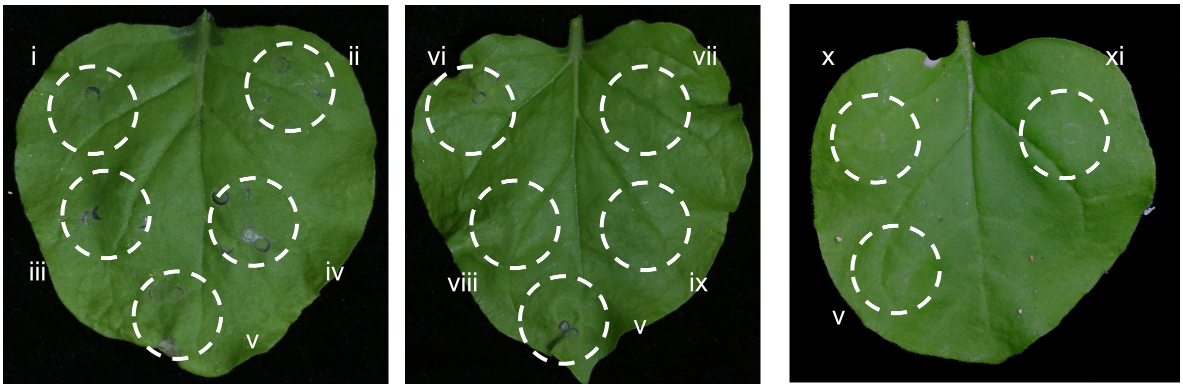


i: 13g184900+SMV P1; ii: 13g184900+SMV Hcpro; iii: 13g184900+SMV 6K1; iv: 13g184900+SMV CI; v: 13g184900+EV; vi: 13g184900+SMV 6K2; vii: 13g184900+SMV VPg; viii: 13g184900+SMV NIa-pro; ix: 13g184900+SMV NIb; x: 13g184900+SMV CP; xi: 13g184900+SMV P3N-PIPO

Supplementary Fig. 3

a

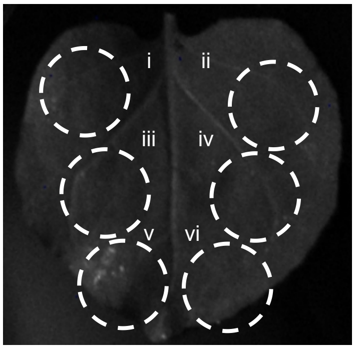


i: NB-cLuc+SMV P3-nLuc;
ii: NB-nLuc+SMV P3-cLuc;
iii: NB-cLuc+EV; iv: SMV P3-cLuc+EV;
v: NB-nLuc+EV; vi: SMV P3-nLuc+EV;

b

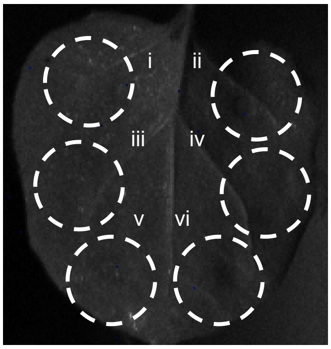


i: LRR-cLuc+SMV P3-nLuc;
ii: LRR-nLuc+SMV P3-cLuc;
iii: LRR-cLuc+EV; iv: SMV P3-cLuc+EV;
v: LRR-nLuc+EV; vi: SMV P3-nLuc+EV;

c

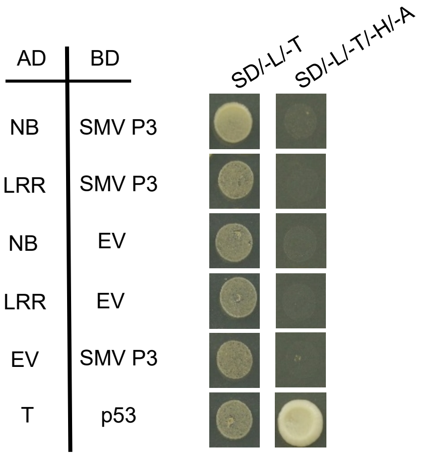


Supplementary Fig. 4

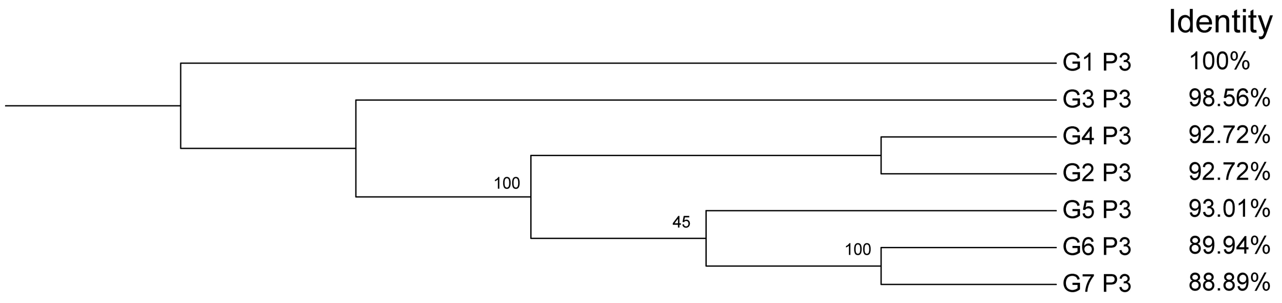


Supplementary Fig. 5

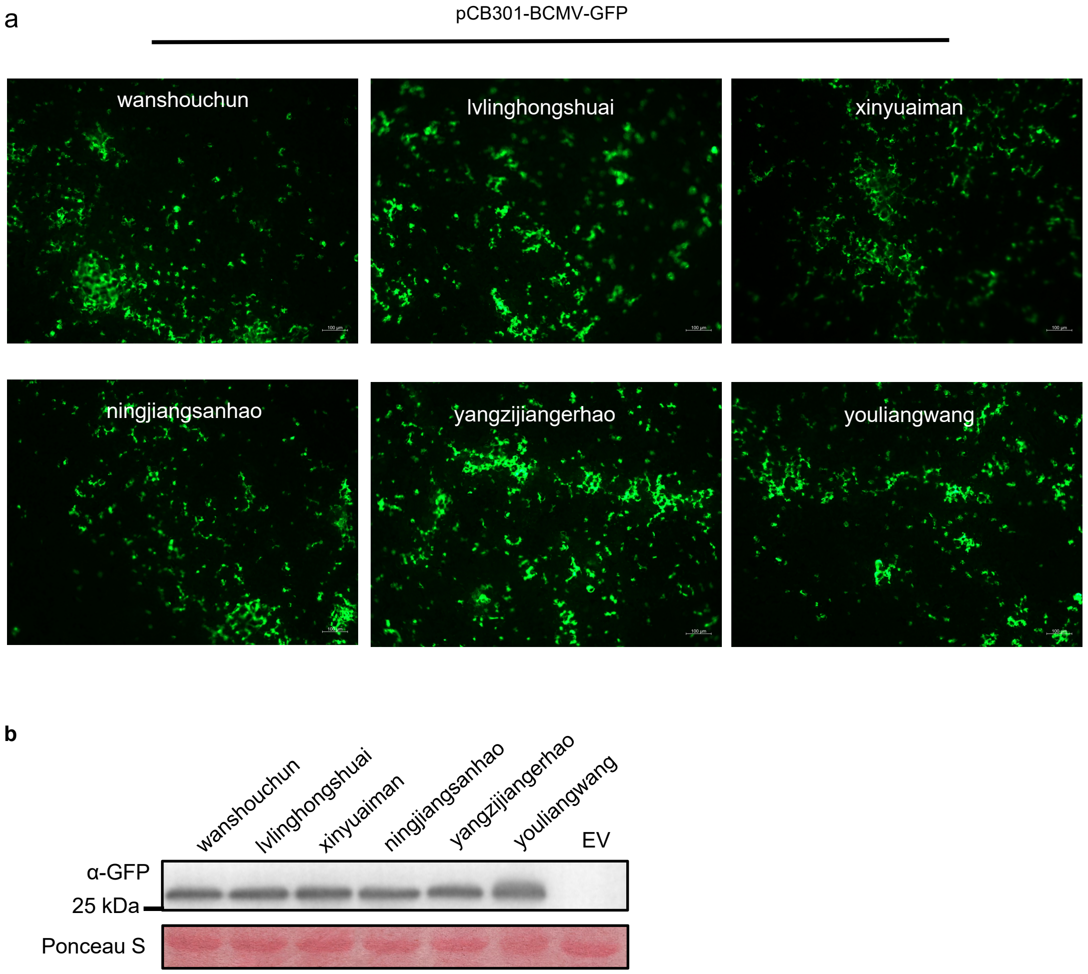


Supplementary Fig. 6

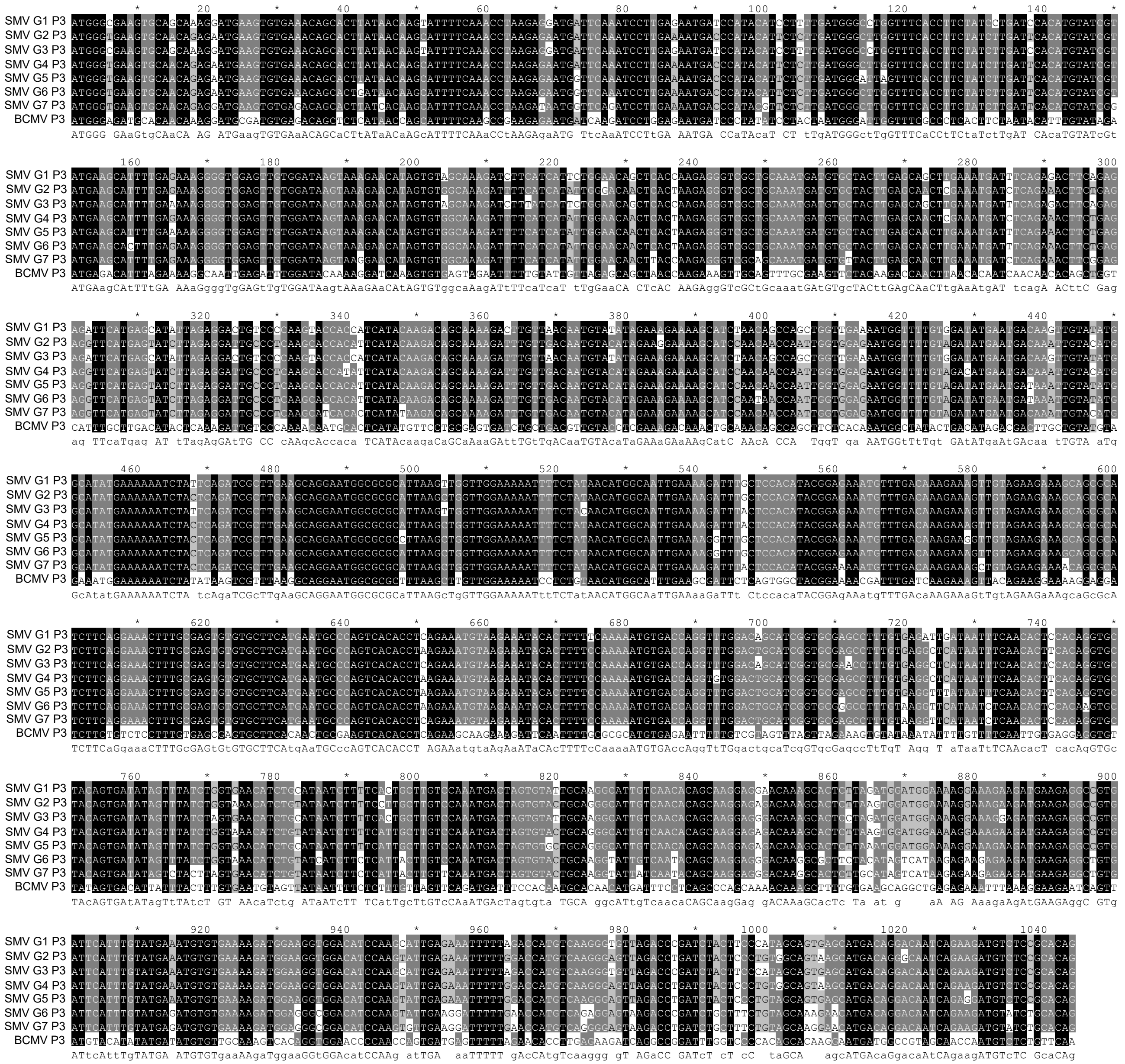


Supplementary Fig. 7
